# Blocking the ability of huntingtin to bind membranes: a therapeutic strategy for Huntington’s disease

**DOI:** 10.1101/2024.07.17.603089

**Authors:** Chathuranga Siriwardhana, Adewale Adegbuyiro, Faezeh Sedighi, Alyssa R. Stonebraker, Sharon Leonard, Maryssa Beasley, Adam Skeens, Blake Mertz, Werner Geldenhuys, Justin Legleiter

## Abstract

The ordered aggregation of proteins into amyloid fibrils is a hallmark of numerous neurodegenerative diseases. A common strategy in developing therapeutics for amyloid-based diseases relies on preventing or manipulating the aggregation process. However, many amyloid-forming proteins and their aggregates bind and damage organelle and cellular membranes. As such, blocking the ability of these proteins from directly interacting with membranes represents a unique therapeutic strategy. Using a mutant huntingtin (htt) protein associated with Huntington’s disease (HD) as a model system, the viability of this strategy was evaluated. Screening over 1200 compounds for their ability to block htt binding to lipid vesicles, two compounds, Ro90-7501 (Ro) and Benzamil (Ben), were identified and validated. Despite directly interacting with htt, neither compound prevented fibril formation. Molecular dynamics simulations suggested each compound has a unique mechanism of action, consistent with experimental data. Importantly, both compounds ameliorated phenotype in a *C. elegans* model of HD.

## Introduction

The aggregation of proteins into amyloid deposits is a hallmark of many neurodegenerative diseases including Alzheimer’s disease (AD), Parkinson’s disease (PD), and Huntington’s disease (HD)^1^. The formation of amyloid fibrils is complex and includes a variety of intermediate oligomers and protofibrils. Many of these different aggregate forms are linked to toxic gains of function, leading to immense effort to develop therapeutic strategies that inhibit or manipulate amyloid formation^2^. However, this approach has had limited success in leading to the development of efficacious drugs. Many amyloid-forming proteins and their aggregates directly bind and damage lipid membranes^3^. For example, high molecular weight oligomers of β-amyloid (Aβ) associated with AD disturb plasma membrane integrity, resulting in decreased membrane fluidity, intracellular calcium dysregulation, depolarization and impaired long-term potentiation^4^. α-synuclein (α-syn) interacts with multiple organelle membranes causing defects in mitochondrial function and membrane trafficking in PD^5,6^. This suggests that blocking the interaction between disease-related proteins (and their aggregate forms) and (sub)cellular membranes is a viable target for therapeutic intervention.

HD is a genetic neurodegenerative disease that is caused by the abnormal expansion of a polyglutamine (polyQ) tract near the N -terminus of the huntingtin protein (htt). PolyQ expansion beyond a critical threshold (∼35 repeats) results in HD. Like other amyloids, htt strongly interacts with various membranous surfaces, and a variety of membrane abnormalities are associated with the presence of mutant htt. Inclusions of htt damage and distort the nuclear envelope in a manner associated with cell death^7^. Furthermore, mutant htt exasperates age-dependent disruption of the nuclear envelope, invoking DNA damage^8^. Htt fibrils impinge on cellular endomembranes, damaging their integrity and freezing ER dynamics^9^. In mammalian cell and primary neuron models of HD, cytoplasmic htt inclusions contain remnants of organelle membranes from the ER and mitochondria, which correlates with impaired organelle function and localization^10^. Damaged organelle membranes (swollen mitochondria and absent nuclear membrane) are also observed in a novel pig model of HD^11^.

Directly preceding the polyQ domain are the first 17 N-terminal amino acids (Nt17, MATLEKLMKAFESLKSF) of htt that acts as a lipid-binding domain^12,13^. In bulk solution, Nt17 is intrinsically disordered but forms an amphipathic α-helix (AH) to facilitate membrane binding^13^. Beyond its role in lipid-binding, Nt17 forms intermolecular associations to form α-helical rich oligomers that promote nucleation of fibril formation^14,15^. This dual role of Nt17 in lipid binding and aggregation is underscored by the direct influence of lipid membranes on the kinetics and mechanism of htt aggregation^14^. This impact of membranes on htt aggregation is highly dependent on lipid composition. For example, total brain lipid extract (TBLE) stabilizes oligomers of truncated polyQ peptides that contain Nt17^12^ and impedes fibrillization of htt-exon1^16^. In contrast, POPC/POPS lipid vesicles enhance aggregation via a unique Nt17-mediated mechanism involving membrane anchoring and two-dimensional diffusion^14^. In general, anionic headgroups accelerate fibril formation^17^ and there is a complementarity relationship between hydrophobic residues and membrane defects, regulating the partitioning of Nt17 into bilayers^18^. The ability of htt to bind membranes is also dependent on aggregation state, with oligomers having the strongest affinity for lipid vesicles in comparison to monomers and fibrils ^19^.

To determine if blocking the ability of amyloid-forming proteins to bind membranes would have potential therapeutic benefit, the interaction of mutant htt with membranes was used as a model system. A systematic screen was performed to identify compounds that block the binding of htt oligomers to lipid vesicles comprised of TBLE. After validation by additional methods, two compounds were identified. The impact of these compounds on htt aggregation and the mechanism by which these compounds block the htt/lipid interaction were then determined. Finally, the ability of these compounds to improve phenotype in a *C. elegans* model of HD was demonstrated. Collectively, our work demonstrates the proof of principle that blocking the binding of amyloid-forming proteins to membranes is a viable therapeutic strategy and provides a method for compound identification and validation.

## Results

### Identifying compounds that block htt/lipid binding

To determine if blocking the ability of htt to bind membranes has therapeutic potential, identification of molecules that directly interfere with the binding process was required. To accomplish this, polydiacetylene (PDA)/TBLE lipid binding assays were used to screen the LOPAC1280 library (Figure 1a). As protein binds to PDA/TBLE vesicles, the associated mechanical stress on the PDA component causes a quantifiable colorimetric response (CR) from blue to red. As htt oligomers have the highest affinity for lipid membranes^19^, PDA/TBLE vesicles were exposed to pre-aggregated, oligomeric htt-exon1(46Q) (10 µM), and the dose of each screened compound was 10x that of htt. As designed, the assay only identified molecules that block htt/lipid interactions by targeting the protein, as compounds that directly associate with PDA vesicles invoke a CR^16^. After 5 h of exposure to htt oligomers, compounds that reduced the CR to 20% of the control response (htt alone) were considered potential hits (Figure 1b). After the initial screen, hits were re-tested in triplicate, and two promising compounds were identified, Ro90-7501 (Ro) and Benzamil (Ben). Ro is a known inhibitor of Aβ and α-syn fibrillization^20,21^. Ben is a potent blocker of the ENaC channel^22^ and a sodium-calcium exchange blocker^23^.

**Figure 1.**
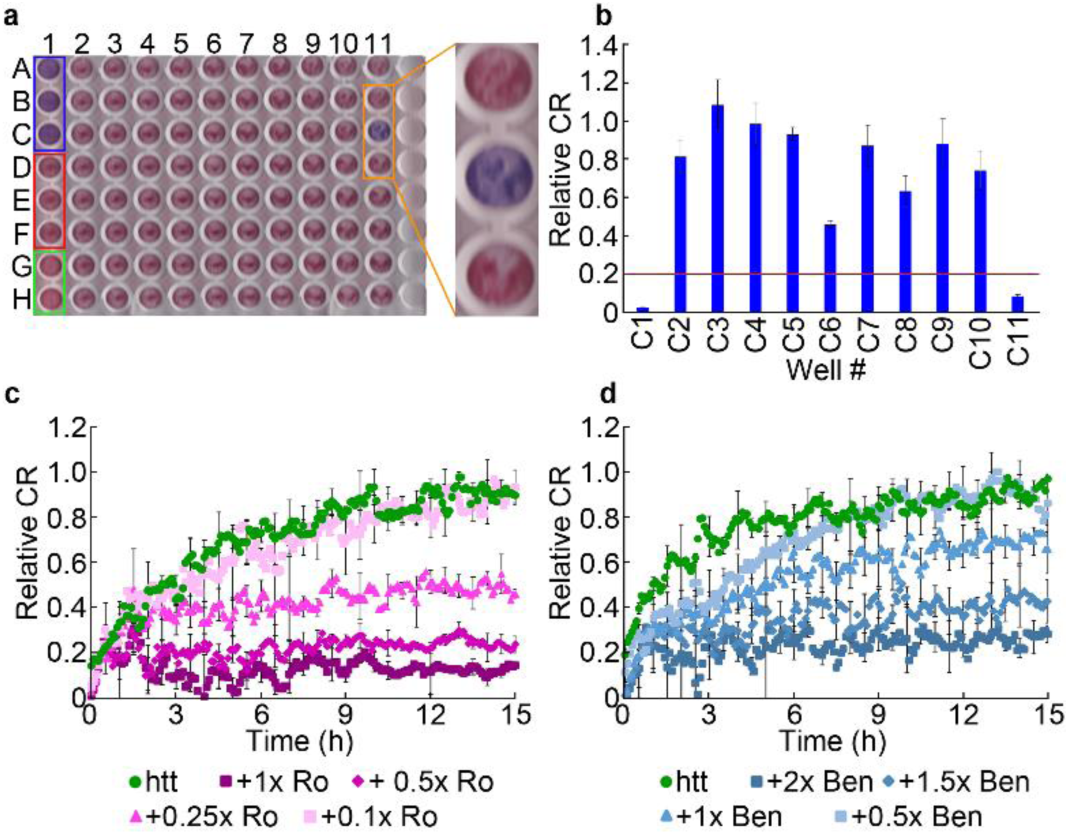
Screen for compounds that inhibit htt binding to lipid membranes. **a,** Representative 96-well plate used for the PDA/TBLE assay after 5 h of exposure to htt-exon146Q. Blue box indicates wells in which vesicles were exposed to neat buffer (negative control). Red box indicates control wells in which vesicles exposed to htt-exon1(46Q). Green box indicated wells in which vesicles were exposed to NaOH (positive control). Inset (indicated by yellow box) highlights wells in which the screened compounds did not inhibit binding to vesicles (top and bottom wells) and a well in which the screened compound did (middle well). **b,** Relative colorimetric response of PDA/TBLE vesicles measured from row C of the plate shown in a. Red line represents the hit threshold. The data is normalized with respect to the htt-exon1(46Q) controls. **c** and **d,** Relative colorimetric response of PDA/TBLE vesicles exposed to htt-exon1 (10 µM) measured kinetically in the presence of varying doses of Ro and Ben normalized with respect to htt control. All error bars represent SEM.

To determine appropriate dosing of each drug for additional validation and experimentation, a series of kinetic PDA/TBLE assays were performed in which the dose of each compound was systematically lowered and the assay time was extended to 15 h (Figure 1c-d). Ro continued to reduce the relative CR below 0.2 with doses as low as 0.5x that of htt. After that, the effectiveness of Ro decreases in a dose-dependent manner until non-significant reduction in relative CR was observed at 0.1x Ro. For Ben, higher doses were needed to suppress vesicle binding. While the 2x dose of Ben reduced the relative CR to ∼ 0.2, a dose of 0.5x Ben no longer provided a statistically significant reduction.

To further validate that the compounds blocked htt/lipid binding, additional assays were performed (Figure 2). First, a calcein dye leakage assay (using POPC vesicles) was performed with effective doses established by the kinetic PDA assays (Figure 2a). Both compounds significantly suppressed dye leakage (p<0.01). With a 1x dose of Ro, the fluorescent signal associated with the dye leakage was reduced to 0.1 relative to the leakage caused by htt. A 2x dose of Ben reduced the leakage to less than 0.2. Next, the impact of htt exposure on supported TBLE bilayer morphology was directly monitored by in solution AFM (Figure 2b). Supported bilayers were formed on mica via vesicle fusion. These bilayers are stable for several hours and have a smooth appearance with an RMS roughness < 0.2 nm. Upon exposure to 10 µM htt oligomers, regions of roughened morphology (RMS of 0.6-1.2 nm) developed on the bilayer surfaces, and the area of membrane disruption grew with time until ∼25% of the bilayer area is altered (Figure 2b-c). When either 10 μM Ro (1x) or 20 μM Ben (2x) was added prior to exposure to htt oligomers, bilayer morphology was relatively unaffected (< 5% of the bilayer area) except for the appearance of a few discrete aggregate structures (Figure 2b-c). Collectively, the PDA, calcein dye, and AFM assays validate that Ro and Ben reduce the ability of htt to bind and damage lipid membranes.

**Figure 2.**
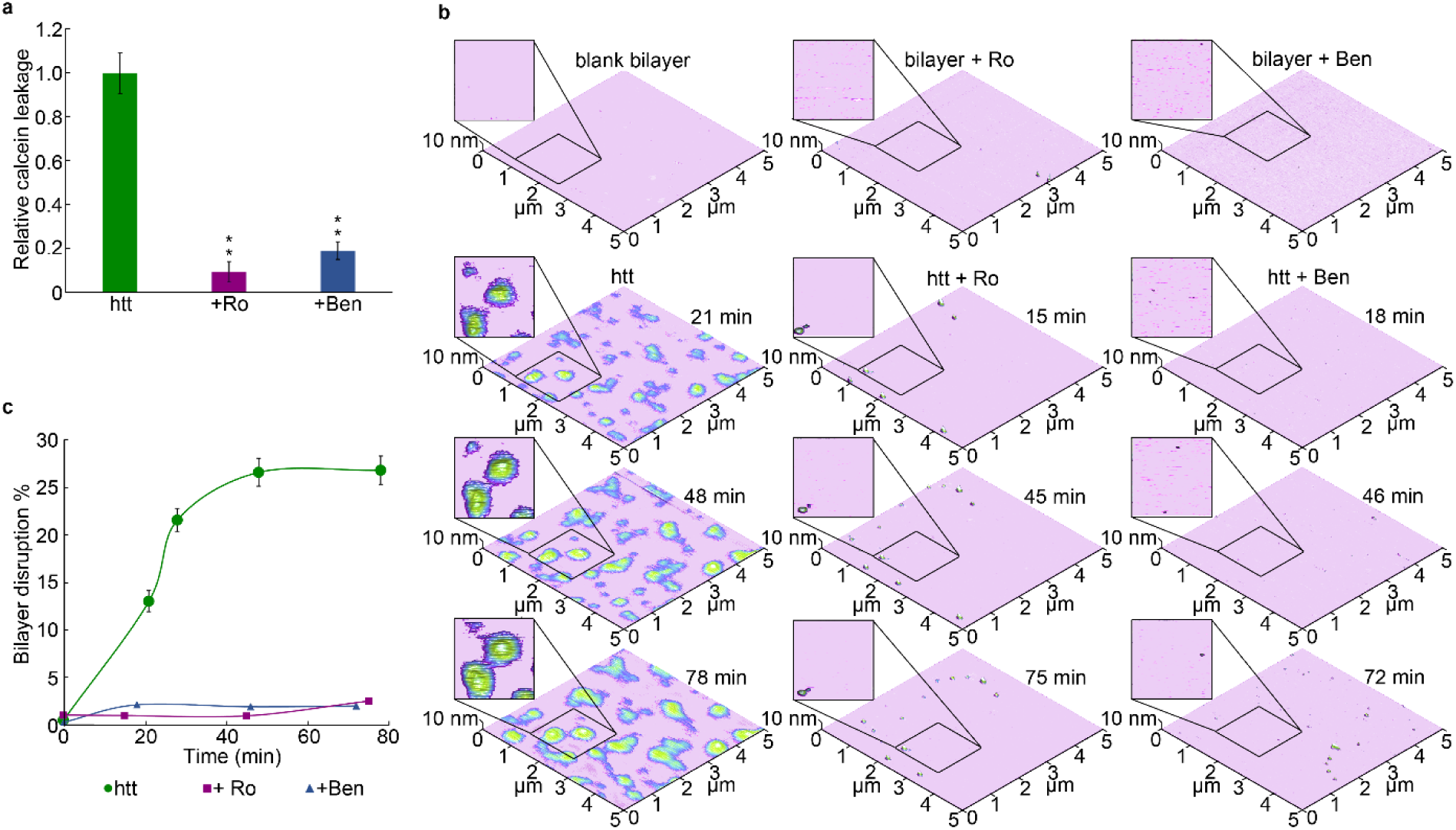
Validation that Ro and Ben inhibit htt binding to lipid membranes. **a,** Calcein dye leakage assay from POPC vesicles exposed to htt-exon1(46Q) (10 µM). Dosing of Ro and Ben were1x and 2x the htt concentration respectively. The data was normalized with respect to htt control. **b,** *In situ* AFM images of supported TBLE bilayers exposed to htt-exon1(46Q) (10µM) or htt-exon1(46Q) incubated with either 1x Ro or 2x Ben **c,** The % area of the bilayer disrupted by exposure to htt-exon1(46Q) determined from *in situ* AFM images. All error bars represent SEM and * indicates P<0.01.

### Impact of Ro and Ben on htt aggregation

As the initial screens were designed to identify compounds that directly interact with htt, it is plausible that Ro and Ben could impact htt aggregation directly. To initially determine if Ro and Ben modify fibril formation, ThT assays were performed using doses of the compounds that effectively inhibited membrane binding. Ro reduced the ThT signal associated with 10 µM htt aggregation in a dose-dependent manner (Figure 3a). However, due to Ro being structurally similar to ThT, there is a possibility that it competes with ThT binding sites on htt fibrils, resulting in reduced ThT signal without inhibiting aggregation. Therefore, htt was pre-aggregated to obtain solutions predominately comprised of fibrils. These solutions were exposed to ThT or a combination of ThT and Ro at the same ratios as the kinetic ThT assays, and the resulting ThT fluorescence was measured (Figure 3b). The ThT signal was reduced in the presence of Ro to comparable levels as observed in the kinetic assays (∼35-40% reduction with 1x Ro). Based on the absorption spectra of Ro (Extended Data Figure 1), it does not interfere with the ThT excitation or emissions signals either. This suggests that Ro competitively bound fibrils and does not suppress htt fibril formation. Ben did not modify fibril formation based on the ThT assay with doses as high as 2x (Figure 3c).

**Figure 3.**
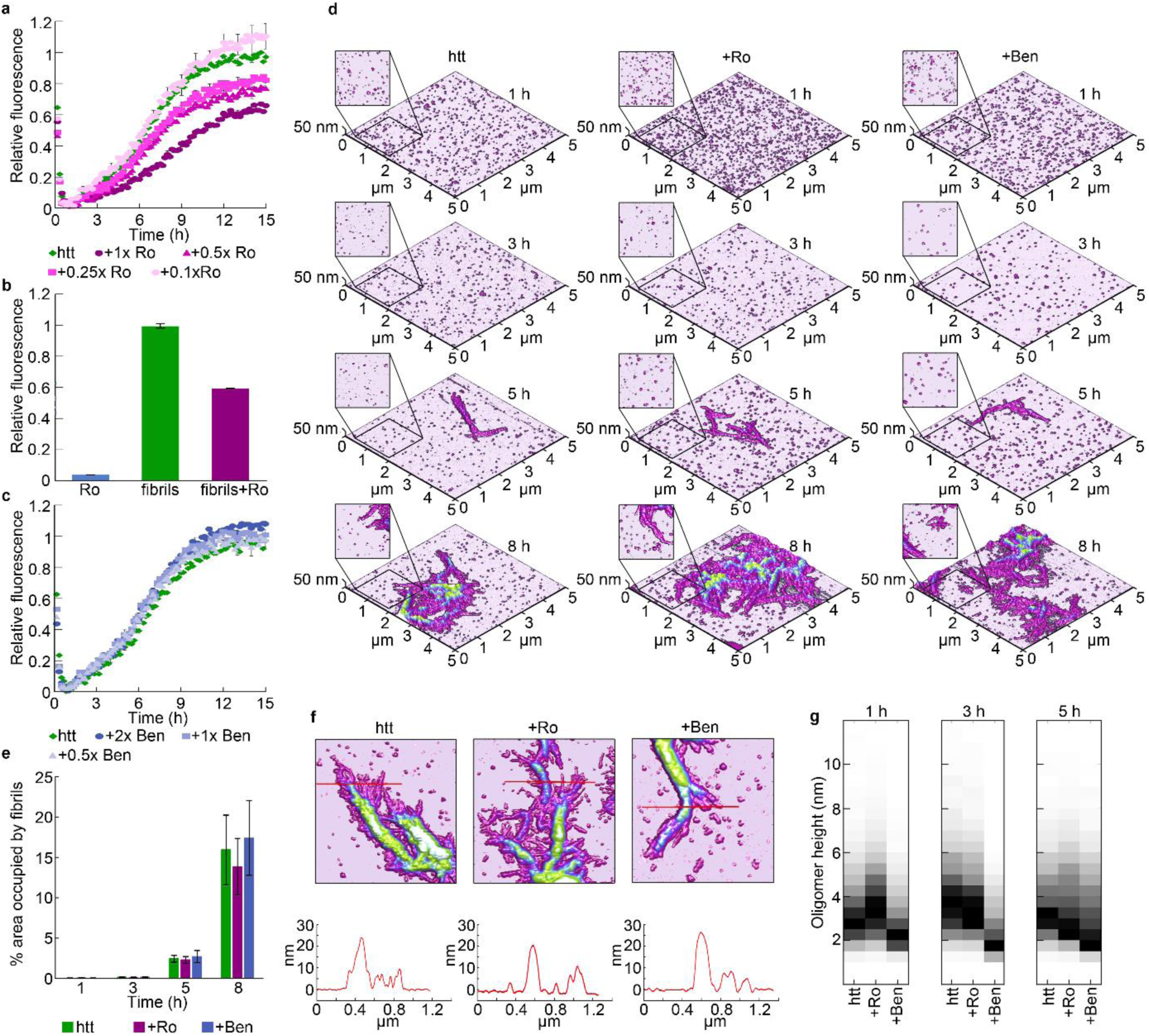
Impact of Ro and Ben on htt aggregation. **a,** ThT fluorescence assay of htt-exon1(46Q) (10 µM) with varying doses of Ro normalized to the htt control. **b,** Relative ThT fluorescence of pre-formed htt-exon1(46Q) fibrils (10 µM) incubated with ThT and ThT +1x Ro. Ro fluorescence without fibrils was used as a control. Signals were normalized to ThT fluorescence of htt fibrils alone. **c,** ThT fluorescence assay of htt-exon1(46Q) (10 µM) exposed to varying doses of Ben normalized with respect to the htt control. **d,** Representative *ex situ* AFM images of htt-exon1(46Q) (10 µM) incubated alone and with 1x Ro or 2x Ben and deposited on mica at various timepoints. **e**, The surface area covered by fibrils calculated from AFM images. **f,** Comparison of fibrils formed in the absence or presence of either 1x Ro or 2x Ben. Red lines in the images correspond to the height profiles directly below the image. **h,** Height histograms of all oligomers formed by htt-exon1(46Q) in the absence or presence of either 1x Ro or 2x Ben as function of time, each column is normalized by dividing the entire distribution by the number of oligomers contained in the most populated bin. All error bars represent SEM.

In addition to the competitive binding of Ro, ThT assays do not provide direct information on potential changes in fibril morphology or oligomer formation. To identify whether Ro and Ben alter morphology of htt aggregates, incubations of 10 µM htt alone, with 1x Ro, or 2x Ben were sampled at several time points and evaluated using *ex situ* AFM (Figure 3d). For all three conditions, only oligomers were observed after 1 and 3 h of incubation, and fibrils emerged after 5 h. At 8 h, all conditions exhibited extensive fibril formation and bundling. Based on the surface area occupied by fibrils at each timepoint, neither compound inhibited fibril formation (Figure 3e), consistent with the ThT assays when taking into account the competitive binding of Ro for htt fibrils. Additionally, fibril morphology was unaltered by the presence of either Ro or Ben, as single fibril strands were typically 7-9 nm thick (Figure 3f). Fibril bundles were also similar in thickness. To characterize oligomer morphology additional image analysis was performed using automated algorithms that measure morphological features of all aggregates present in an AFM image (Figure 3g). In this analysis, oligomers were defined as objects greater than 0.8 nm in height and with an aspect ratio less than 2.5 (globular structure) and occupying a surface area < 0.008 µm^2^. After 1, 3, and 5 h of incubation, oligomers formed by htt alone or with Ro were similar in size (mode height ∼2.5-4 nm,); however, the addition of Ben resulted in systematically smaller oligomers (mode height ∼1.5 – 2.5 nm), suggesting that Ben altered oligomer structure without reducing their efficiency in nucleating fibrilization.

### Mechanism of interaction of Ro and Ben with htt

As Nt17 facilitates the interaction of htt-exon1 with lipid membranes^12,13^, it is likely that Ro and Ben directly interact with Nt17. To gain insight into this potential interaction, molecular dynamics (MD) simulations were performed with both monomeric and oligomeric forms of Nt17 exposed to each compound (Figures 4-5 and Extended Data Figures 2-5). For the oligomer structure, the tetramer of Nt17 resolved by Kotler et al was used^24^. This structure is a dimer of dimers having anti-parallel Nt17 helical units. It is important to point out that oligomers observed by AFM in this study are typically larger than tetramers; however, this NMR structure is the only one currently available. For the monomeric simulations, a single Nt17 peptide from the tetramer structure was used and was therefore initially α-helical. The NMR structure was determined from peptides lacking the first methionine residue of Nt17 because this residue is typically cleaved off *in vivo*. For ease of comparison later with htt-exon1 experiments, the amino acids are numbered here with the missing methionine being considered the first residue.

**Figure 4.**
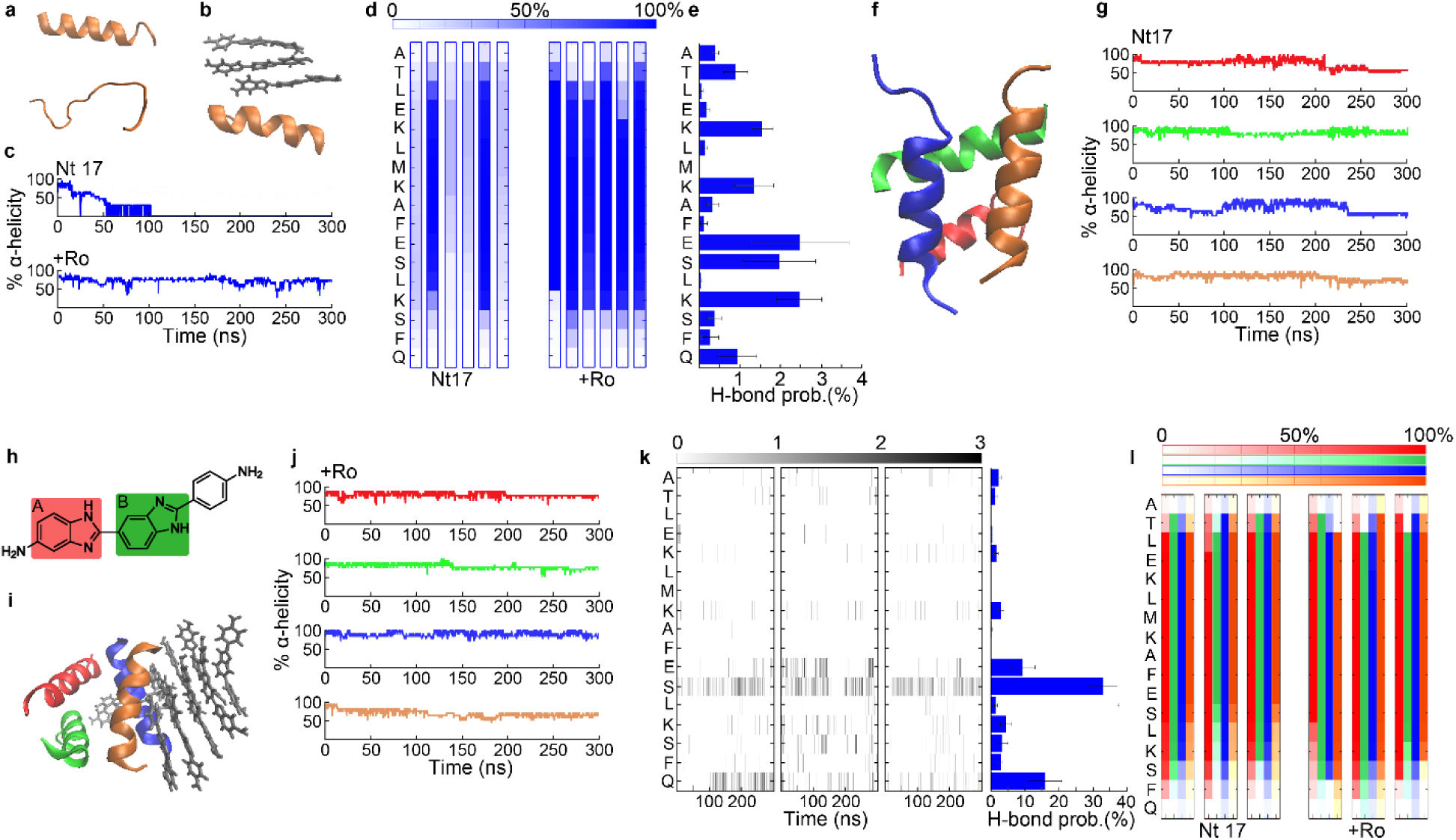
MD simulations of the interaction of Ro with Nt17. **a,** The initial Nt17 monomer structure and after 150 ns of simulation. **b,** Nt17 monomer interacting with Ro molecules after 150 ns of simulation. **c,** The percent α-helicity of Nt17 monomer in the presence and absence of Ro as a function of time (five additional simulations provided in Extended Data Figure 2). **d,** The average percent α-helicity of each residue in Nt17 monomers in the absence and presence of Ro. For each condition, the individual heat maps represent an independent simulation. **e,** Residue specific, hydrogen bonding probability density of Ro with Nt17 monomer averaged across six independent simulations. **f,** Nt17 tetramer structure after 150 ns of simulation. **g,** The percent α- helicity of each Nt17 peptide within the tetramer structure as a function of time (two additional simulations provided in Extended Data Figure 4). **h,** Structure of Ro with each benzimidazole highlighted. **i**, Nt17 tetramer structure exposed to Ro molecules after 150 ns of simulation. **j,** The percent α-helicity of each Nt17 peptide within the tetramer structure upon exposure to Ro as a function of time(three additional simulations provided in Extended Data Figure 4). **k,** Residue specific, time-dependent hydrogen bonding profile and probability density of Ro with the Nt17 tetramer. Each heat map represents an independent simulation. **l**, The average percent α-helicity of each residue in Nt17 broken down by peptide within the tetramer structure in the absence or presence of Ro. Three independent simulations are shown for each condition. Each column consists of 4 heat-maps representing the different Nt17 peptides (color coded). All error bars represent SEM.

**Figure 5.**
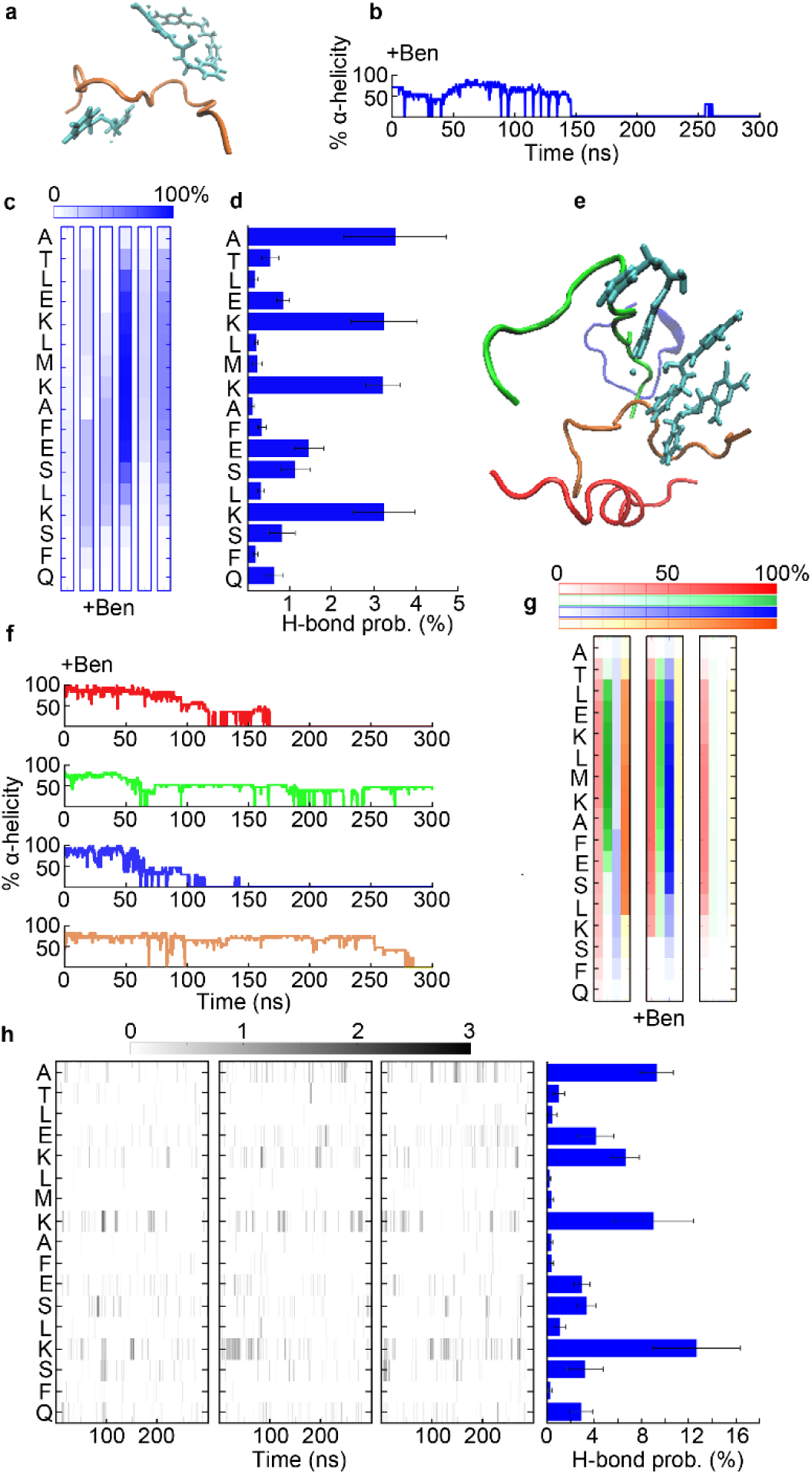
MD simulations of the interaction of Ben with Nt17. **a,** Nt17 monomer structure exposed to Ben after 150 ns of simulation. Nt17 is represented in orange and Ben in cyan. **b,** The percent α-helicity of Nt17 monomer in the presence of Ben as a function of time (five additional simulations provided in Extended Data Figure 2). **c,** The average percent α-helicity of each residue in Nt17 monomers in the presence of Ben. For each condition, the individual heat maps represent an independent simulation. **d,** Residue specific hydrogen bonding probability density of Ben with Nt17 monomer averaged over six independent simulations., time-dependent hydrogen bonding profile and probability density of Ben with an Nt17 monomer for three independent simulations. **e,** Nt17 tetramer structure exposed to Ben molecules after 150 ns of simulation. **f,** The percent α-helicity of each Nt17 peptide within the tetramer structure upon exposure to Ben as a function of time (two additional simulations provided in Extended Data Figure 4). **g,** The average percent α-helicity of each residue in Nt17 broken down by peptide within the tetramer structure in the presence of Ben. Three independent simulations are shown for each condition. Each column consists of 4 heat-maps representing the different Nt17 peptides (color coded). **h,** Residue specific, time-dependent hydrogen bonding profile and probability density of Ben with the Nt17 tetramer for three independent simulations. The color bar represents the number of H-bonds formed. All error bars represent SEM.

While the initial structure of the Nt17 monomer was helical, this secondary structure was unstable in the absence of any compounds and became completely disordered within 50-100 ns in four out of six simulations (Figure 4a-c and Extended Data Figure 2a). Even in the simulations in which helical structure was present for the vast majority of the simulation time, there were several instances in which the Nt17 completely unfolded and refolded (Extended Data Figure 2a). Addition of three Ro molecules stabilized the α-helical structure of Nt17 (Figure 4b-d and Extended Data Figure 2b). Comparison of the α-helical probability of each residue in Nt17 in the absence and presence of RO across each simulation demonstrates a clear stabilization of the helical structure (Figure 4d). While Ro would intermittently form H-bonds with most residues of Nt17 (M8 being an exception), these H-bonds were short-lived (typically in the range of 0.3±0.1 ns and occurring less than 5% of the total simulation time for any given residue, Figure 4e and Extended Data Figure 3a). Ro-907501 had a slight preference for interaction with the adjacent residues E12 and S13, as well as K15 (Figure 4e). Binding of additional Ro molecules to an Nt17-Ro complex was facilitated by extensive π-π stacking upon the bound Ro molecule.

When simulated in the absence of small molecules, the Nt17 tetramer was stable but displayed structural flexibility, as the helical content of each peptide would fluctuate to as low as ∼50% helicity (Figure 4g and Extended Data Figure 4a). Addition of ten Ro molecules stabilized the tetra-helical bundle by forming H-bonds with the compound (Figure 4h-j and Extended Data Figure 4b). Residues sequestered within the tetramer structure did not readily form H-bonds with Ro. However, Ro had a strong preference for the C-terminal end of Nt17 peptides in the oligomer, and in particular H-bonded to S13 (∼35% of the simulation time, Figure 4k and Extended Data Figure 5). The H-bonds with S13 would typically form with the NH group of the first benzimidazole (labeled A in Figure 4h). To a lesser extent (∼10% of the time), this NH group formed H-bonds with E12. Once Ro was positioned at S13 via H-bonding, the NH group of the second benzimidazole (labeled B in Figure 4h) and the 5-amine group of benzimidazole intermittently H-bonded with other residues (both intermolecularly and intramolecularly, Extended Data Figure 5) with a preference for C-terminal residues and particularly Q18 (∼15% of the simulation time). Similar to the interaction of Ro with monomers, additional Ro molecules would π-stack upon the Ro bound to Nt17 (Figure 4i). Overall, interaction with Ro stabilized the α-helical structure of the C-terminal end of Nt17 peptides within the tetramer relative to the control (Figure 4l). The α-helix of Nt17 peptides in control simulations readily unfolded from the C-terminal end to E12, and the addition of Ro-90-7501, by associating with S13 and other C-terminal residues, significantly restricted this unfolding. This suggests structural flexibility at the C-terminal side of Nt17 plays a key role in htt oligomers binding membranes, consistent with recent cross-linking studies^19^.

As S13 represented the primary site of association between htt and Ro, additional PDA/TBLE lipid binding assays were performed with a htt-exon1(46Q) construct containing a S13D mutation (Extended Data Figure 6). Adding a negative charge at position 13 should strengthen the association between oligomers and Ro, enhancing the effectiveness of the compound to block htt/lipid interactions. Indeed, Ro reduced the CR associated with exposure to S13D htt-exon1(46Q) below 20% of control with doses as low as 0.1x (an ineffective dose with wild type htt). This was specific to Ro, as the S13D mutation had no impact on the efficiency of Ben to block htt binding to vesicles.

Ben interacted with Nt17 in a manner distinct from Ro (Figure 5a-d and Extended Data Figure 2c). In simulations with monomers, Nt17 completely unfolded within ∼60-150 ns in five out of six simulations in the presence of three Ben molecules. While intermittent H-bonds formed between Ben and each residue within Nt17 (less than 6% of simulation time for all residues), a slight preference for interacting with lysine residues (K6, K9, K15) and A2 due to the presence of the N-terminal amine group (Figure 5d). The lifetime of H-bonds with these amine-containing residues was typically longer on average (∼0.5 nm compared to ∼0.3 ns for other residues, Extended Data Figure 3b). Overall, Ben appeared to have a minimal impact on the behavior of Nt17 monomers compared with control.

With Nt17 tetramers, the addition of Ben greatly destabilized the α-helical bundle structure (Figure 5e-g and Extended Data Figure 4c); at least two or more α-helices were completely unfolded in each simulation. Nt17 peptides that were not completely unfolded by Ben displayed considerably less helicity compared with the control (Figure 5g). This unfolding allowed Ben to form intermittent H-bonds with all residues, and the preference for the N-terminal amine group and the three lysine residues remained (Figure 5h). However, H-bond formation with K15 was enhanced. K6 and K15 participate in salt bridges that stabilize htt oligomers^25^, and Ben appears to interfere with these stabilizing interactions. Interestingly, this reduced stability of the tetramer structures with minimal impact on Nt17 monomers is consistent with the smaller oligomers observed via AFM (control oligomer mode height of 4-5 nm compared with oligomer mode height of 1.5-2.5 nm in the presence of Ben). As the development of helical structure in Nt17 is associated with lipid binding, the ability of Ben to destabilize the Nt17 α-helix would reduce the ability of htt to bind membranes.

### Ro and Ben rescue phenotype in a C. elegans model of HD

To determine if Ro and Ben could ameliorate HD phenotype, both compounds were tested on a *C. elegans* model of HD, which express htt513 (slightly longer than exon1)^26^. The control strain, EAK102, expresses a polyQ stretch of 15 and will be referred to as Q15; EAK103 has a repeat glutamine stretch of 128 and will be referred to as Q128. Importantly, both are labeled with YFP at the C-terminal end of htt513, limiting the interference of this tag with the ability of Nt17 to bind lipids. Dosing of both compounds was optimized by performing viability assays on WT *C. elegans* to determine the largest, nontoxic dose. Worms were treated with 150 µM of Ro and 150 µM of Ben. In age synchronized populations, the Q128 strain is less viable compared with Q15 as a function of time. Both Ro and Ben significantly (p < 0.05) increased viability of Q128 worms to Q15 levels by day 23 (Figure 6a). Beyond viability, Q128 worms develop a thrashing deficit as early as day 2 of adulthood. Treatment with Ro or Ben significantly improved (p<0.01) thrashing deficit on day 2 of adulthood and restored thrashing rates to control Q15 worm levels by day 7 (Figure 6b). Q128 worms develop visible inclusions of htt by day 2 of adulthood that do not form in Q15 worms. Despite improving viability and rescuing thrashing deficits, neither Ro or Ben prevented the development of these inclusions (Figure 6-d), which is consistent with neither compound impacting fibril formation *in vitro*.

**Fig 6.**
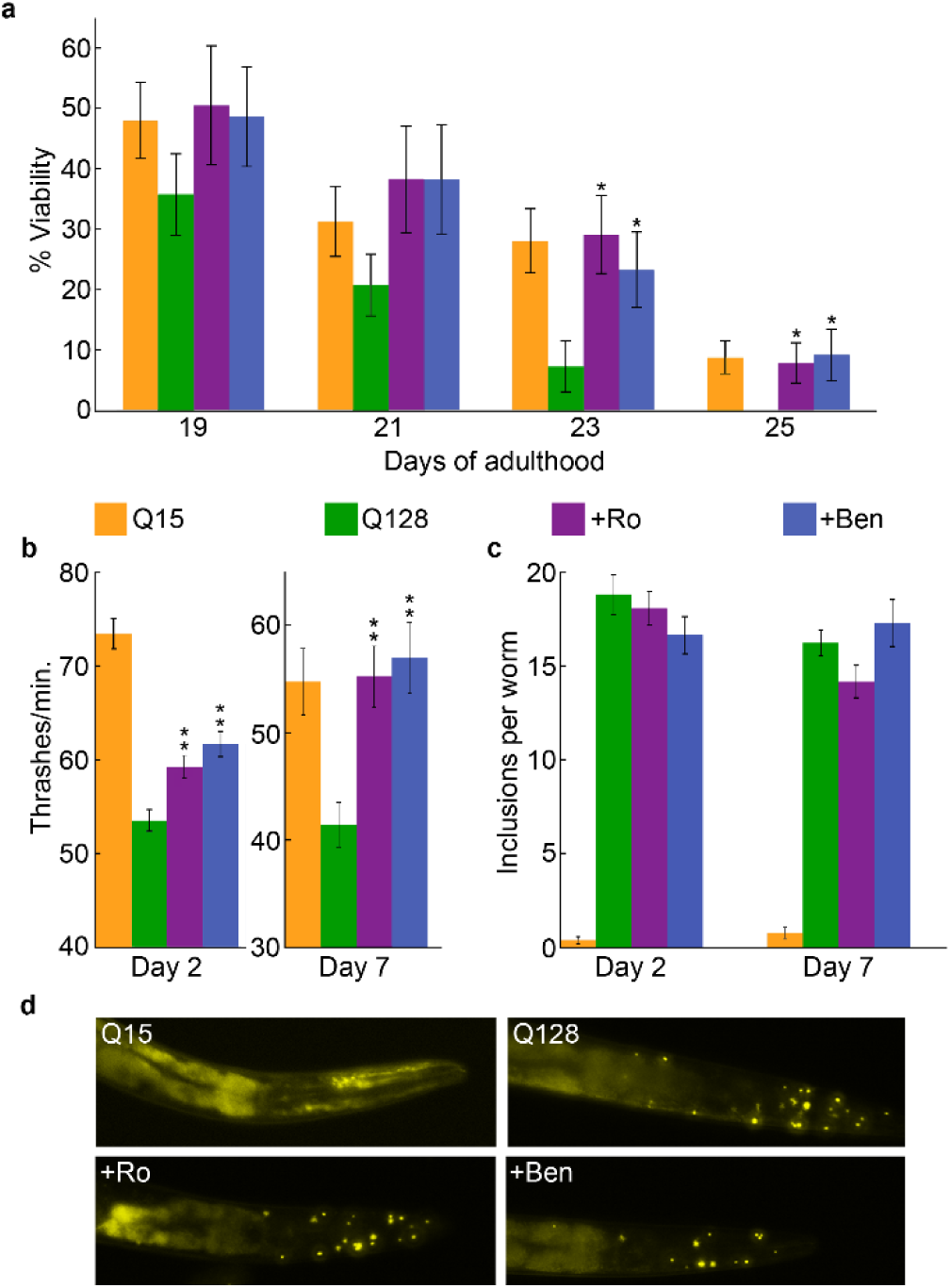
Ro and Ben ameliorate phenotype in a *C.elegans* model of HD. **a,** Viability of Q15 and Q128 worms. Q128 worms were treated with either Ro or Ben. **b,** Thrashing rate measured in liquid media of Q128 worms untreated or treated with either Ro or Ben. Q15 worms were used as a control in all experiments. **c,** Number of inclusions formed per worm based on fluorescent microscopic images. **d,** Representative fluorescent images of Q15 worms and Q128 worms either untreated or treated with Ro or Ben. All error bars represent SEM and * indicates P<0.05 and ** indicates P<0.01

## Discussion

Historically, modifying aggregation has been a popular strategy for developing therapeutics for amyloid-based diseases. While numerous methods to alter aggregation have been developed, ranging from small molecules to antibodies^27–30^, this strategy has not yielded an effective therapeutic. Of interest here, many small molecule aggregation inhibitors lose their effectiveness in the presence of membranes^16,31^. Strikingly, many amyloid proteins readily associate with and damage a variety of cellular membranes. Here, a novel strategy was developed that targets the ability of amyloid-forming proteins from directly interacting with lipid membranes. Using htt-exon1 as a model system, two compounds were identified, their mechanism of action explored, and their ability to rescue phenotype in a *C. elegans* model of HD demonstrated. Collectively, this study serves as a proof of principle for this approach.

Ro has been identified as a potentially generic inhibitor of amyloid formation. In this regard, Ro effectively inhibits both Aβ and α-syn aggregation^20,21^. While ThT data suggests that Ro competitively bound to fibrils, it was not effective in modifying htt aggregation. As htt fibrils exhibit limited ability to bind lipid vesicles and the screens were performed with non-fibrillar htt, the ability to bind fibrils does not appear to be the major factor in blocking htt/lipid interactions. MD simulations suggested that Ro has a high propensity to localize to S13 in Nt17 in both monomeric and oligomeric forms of htt. Additional molecules of Ro can subsequently π-stack on top of the Nt17 bound Ro. This π-stacking may provide shielding between polar residues that facilitate the initial steps in htt monomers binding to membranes^32^. Within oligomers, the Ro molecule bound to S13 forms intermittent hydrogen bonds (both intra-and intermolecular) with other residues toward the C-terminus of Nt17. These contacts stabilize the α-helical structure of Nt17 and reduce the configurational flexibility of the oligomer. Cross-linking studies suggest that configurational flexibility plays an important role in the ability of htt oligomers to bind lipid membranes^19^. The importance of the ability of Ro-90-7501 to localize to S13 is experimentally supported by the enhanced efficacy of blocking htt/lipid interaction of htt containing a S13D mutation.

Ben has been shown to reduce htt-induced toxicity in cellular models and mouse models^33^. Ben, as an acid sensing ion channel blocker, reduced htt toxicity in HD mice by reducing acidosis and enhancing ubiquitin proteosome activity. Our study suggests that Ben also directly interacts with htt, blocking its ability to bind membranes and providing additional protection. The mechanism by which Ben blocks htt from binding membranes differs from that of Ro. Based on MD simulations, Ben uncoils the α-helical oligomer structure by preferentially interacting with amine groups at the N-terminus and in lysine residues of Nt17. Lysine residues in Nt17 are highly protected in aggregate structures^25^, suggesting they are involved in stabilizing interactions such as salt bridges. However, the interaction of Ben with monomers is much weaker compared with oligomers. Collectively, this results in slightly smaller oligomers as the rate of dissociation is slightly increased while the association rate is mostly unaffected. However, a steady population of oligomers develops, resulting in no impact of fibril formation. The destabilization of the underlying α-helical structure may underscore the reduced affinity for lipid membranes. It is also possible that by localizing near Lysine residues that Ben blocks electrostatic interactions between Lysine and anionic headgroups of lipids. This is consistent with acetylation of these lysine residues reducing the affinity of htt for lipids^34^. The apparent disparate modes of action by which Ro and Ben block htt/lipid interactions points to a multi-faceted mechanism by which htt monomers and oligomers bind and disrupt membranes.

Despite improving phenotype in *C. elegans*, neither drug eliminated inclusion formation. As neither drug had an inhibitory impact on fibril formation *in vitro*, this is not surprising. However, the situation *in vivo* is likely more complicated. Lipid membranes have a varied impact on htt aggregation depending on lipid composition. POPC, POPS, DMPC, POPG, and POPC/POPS mixed systems all promote fibril formation^14,17,18^. DOPC, POPE, TBLE, cardiolipin, inner and outer mitochondrial mimics, and other cellular extracts decrease fibril formation^17,18,35,36^. With TBLE and other cellular extracts decreasing aggregation, htt association with lipid membranes *in vivo* may slow aggregation into inclusions. As such, blocking htt from those membranes may enhance inclusion formation. This notion that impeding the ability of htt to bind membranes would enhance aggregation is supported by the observation that removing Nt17 from htt-exon1 expressed in cells enhances inclusion formation and the efficiency of seeding aggregation with pre-formed aggregates^37^. Additionally, crowded aqueous environments (similar to the cytosol) at membrane/liquid interfaces further modifies htt aggregation on membranes in a manner dependent on crowding agents^38^.

This approach of blocking the ability of amyloid-forming proteins to directly interact with lipid membranes can be extended toward other amyloid-based disease and biological processes. Lipids are a common modulator of amyloid formation, as studies of α-synuclein^39^, islet amyloid polypeptide^40^, β-amyloid^41^, and polyQ^12,14,42^ all demonstrate altered aggregation in the presence of membranes and the ability to induce physical damage to membranes. This notion is supported by recent reports that the natural products, squalamine and trodusquemine, ameliorate α- synuclein and Aβ toxicity associated with PD and AD respectively by displacing oligomers from membranes^43,44^. With regard to Aβ, squalamine and its derivatives ameliorated Aβ-induced toxicity despite increasing amyloid formation^45^. The mechanism of action of these natural products is via changing membrane properties^44^. Our strategy targets protein aggregates directly, gaining specificity. Additionally, amyloids function in viral and bacterial infection, potentially via membrane interaction. For example, amyloid aggregates of PAP248-286 in human semen severely increase infection efficiency of HIV^46^ as PAP248-286 fibrils promote interaction between the lipid membrane of the HIV virion with the host membrane^47^. Murine cytomegalovirus protein M45^48^, NSs (Rift Valley fever virus)^49^, and the gut bacterial protein curli^50^ are functional amyloids associated with infection. Thus, targeting the ability of amyloids to bind and damage membranes may find broad utility as a therapeutic strategy.

## Supporting information

ExtendedMaterials

## Notes

### Competing Interest Statement

The authors have declared no competing interest.

