## ExtendedMaterials for "Blocking the ability of huntingtin to bind membranes: a therapeutic strategy for Huntington’s disease"

Extended Material for:

### Methods

*Purification of GST-htt-exon1 fusion proteins.* Glutathione *S*-transferase (GST)-htt-exon1 fusion protein with 46 glutamine repeat units was purified as described previously[1]. In brief, GST-htt exon 1 (htt) was expressed in *Escherichia coli* by induction with isopropyl $\beta$-D- thiogalactoside at 30 °C for 4 h. Cells were lysed with a combination of lysozymes (0.5mg/mL) and probe sonication. The protein was purified from the lysate with a GST affinity column using LPLC, Bio-rad liquid chromatography. Relevant fractions were identified using ultraviolet absorption and confirmed by gel electrophoresis. The protein was subjected to dialysis for 48 h. Before use in any assay, the protein was centrifuged for 30 min at 20000 ×g at 4 °C to remove preexisting fibrils. Factor Xa (New England Biolabs) was used to cleave the GST tag. All experiments were carried out in a buffer composed of 50 mM Tris-HCl, 150 mM NaCl.

*TBLE/PDA assay.* TBLE/PDA assays were performed as reported previously[2]. Shortly, TBLE (Avanti lipids) and 10,12 tricosadiynoic acid were combined in a molar ratio of 2:3 in an ethanol/chloroform mixture. Solutions were dried under a stream of $N_2$ for 15 min. The resulting film was re-suspended in 50 mM tris buffered saline at 70 °C. After 5 min of probe sonication, solutions were stored at 4 °C overnight to allow self-assembly of vesicles. The diacetylene monomers were polymerized by irradiation under 254 nm for 15 min, resulting in a bright blue color solution. Prior to use in TBLE/PDA assay, htt-exon1(46Q) was incubated on ice with Factor Xa for 90 minutes to allow cleavage and formation of oligomers. Final concentration of htt-exon1(46Q) in TBLE/PDA assays was 10 µM.

For TBLE/PDA screening assays, the LOPAC 1280 library was used in a 96 well plate format, with each plate screening 80 compounds. Polymerized PDA/TBLE vesicles were added to all the wells in the plate excluding column 12. Column 1 was reserved for controls. As a negative control, 50 mM tris saline buffer was added to cells A1, B1, and C1. Wells D1, E1, and F1 were exposed to htt-exon1(46Q). As a positive control, saturated NaOH was added to wells G1 and H1 to ensure PDA/TBLE vesicles were working properly. A unique compound was added to wells in columns 2-11 at a final concentration of 100 µM, along with 10 µM htt-exon1 making the final compound to htt ratio 10:1 in the screening assays. Plates were incubated at 25 °C and colorimetric response (CR) was determined at 1 h, 3 h, and 5 h by measuring the blue absorbance (640 nm, $A_{blue}$) and the red absorbance (500 nm, $A_{red}$) (Spectramax M2) and using:

$$CR = \frac{PB_0 - PB_1}{PB_0} \quad (1)$$

where PB is defined as $A_{blue}/(A_{blue} + A_{red})$. $PB_0$ is determined from the negative control, and $PB_1$ is obtained from each experimental condition.

For PDA/TBLE kinetic assays, polymerized PDA/TBLE vesicles were exposed to htt-exon1 (46Q) at a final concentration of 10 µM in the presence of either Ro90-7501 or Benzamil (TOCRIS) at various ratios. 50 mM tris saline buffer was used as the negative control. CR was determined every 5 min for 15 h. All experiments were carried out in 96-well plates at 25 °C.

*Calcein dye leakage assay.* Large unilamellar vesicles (LUVs) of POPC with entrapped calcein were prepared as described[3]. Lipid films were suspended in calcein buffer (70 mM calcein 10 mM Tris, 150 mM NaCl pH 7.4). To promote LUV formation, the lipid solutions were subjected to 10 freeze-thaw cycles followed by bath sonication for 30 min. Excess dye was removed with size exclusion chromatography using Sephadex G-50 beads (Sigma-Aldrich) using 10 mM Tris, 150 mM NaCl as the eluent. Vesicles were exposed to 10 µM htt-exon1(46Q) in the absence and presence of Ro (1x) and Benzamil (2x). Calcein fluorescence was measured at 515nm (495 nm excitation) prior to the addition of protein and 10 min after the addition of protein at 37 °C. Vesicles were then lysed with 1% Triton X-100 to obtain maximum fluorescence. Relative leakage was calculated by

$$Relative\ leakage = \frac{I_f - I_i}{I_t - I_i} \qquad (2)$$

where $I_f$ is the fluorescent intensity after addition of protein, $I_i$ is the fluorescent intensity before addition of protein, and $I_t$ is the fluorescent intensity after lysis.

*Atomic force microscopy (AFM).* All AFM imaging was performed with a Nanoscope V Multimode atomic force microscope (Veeco, Santa Barbara, CA) equipped with a closed-loop vertical engage J-scanner. For *ex situ* AFM experiments to determine the impact of compounds on aggregation, htt-exon1(46Q) (10µM) was incubated in the absence and presence of Ro or Benzamil at 37 ˚C and sampled after 1 h, 3 h, 5 h and 8h for imaging. At each time point, a 5 µL aliquot of each sample was deposited on freshly cleaved mica, allowed to set for 1 min, washed with ultrapure water, and dried with a stream of air. Imaging was done using silicon oxide cantilevers with a normal spring constant of ~40 N/m and resonance frequency of ~300 kHz at a scan rate of ~2Hz.

For *in situ* AFM imaging of htt-exon1(46Q) occurring directly on lipid bilayers, silicon nitride cantilevers with a spring constant of ~0.1 N/m were used. 25 µL aliquots of TBLE vesicles were injected directly into the fluid cell at a concentration of 1 mg/mL. The formation of a continuous defect free bilayer on mica via vesicle fusion was observed by continual AFM imaging. Htt-exon1(46Q) that had been incubated at 4 °C for 90 min with factor Xa was injected into the fluid cell to a final concentration of 10 µM. For experiments with either Ro (2x) or Benzamil (1x), the drug was incubated with the htt-exon1(46Q) prior to injection into the fluid cell. Images obtained by both *ex situ* and *in situ* AFM were processed and analyzed using MATLAB as described previously[4].

*Thioflavin T (ThT) assay.* ThT assays were conducted in 96- well plates on a Spectramax M2 microplate reader (Molecular Devices). Final ThT concentration in each well was 40 µg/mL. The aggregation of htt in the presence of varying doses of Ro or Benzamil was investigated by measuring ThT fluorescence at 484 nm emission (440 nm excitation) at 37 °C for 15 h with readings taken every 10 min. All experimental conditions were performed in triplicates. For analysis of Ro's ability to compete with ThT to bind fibrils, htt-exon1(46Q) was incubated at 37 °C for 24 h to obtain a predominately fibrillar sample. Fibrils were then incubated with either ThT or ThT with Ro ensuring a final htt:Ro ratio of 1:1 at 37 °C for 1 h and ThT fluorescence was measured.

*Molecular dynamic (MD) simulations.* A Nt17 tetramer structure by Kotler et al (6N8C) was utilized to perform MD simulation[5]. For monomer simulations one peptide from the tetramer structure was extracted. Initially the peptides were solvated in water (TIP3) and neutralized with NaCl (150 mM) using Charmm-gui[6] and was subjected to energy minimization and equilibration. Ro and Benzamil were then introduced to the equilibrated peptide structures using packmol[7]. For monomer and tetramer simulations, three and ten drug molecules were added respectively. The resulting structures (peptide + drug) were then solvated and ionized to a final NaCl concentration of 150 mM using VMD solvate and ionize plugins[8]. Ro and Benzamil were parameterized with CHARMM General Force Field (CGenFF)[9]. Nano-scale molecular dynamics (NAMD 2.14)[10] was used to perform all simulations on CHARMMc36 force field[9]. The temperature and pressure parameters were set to 310 K and 1.01325 bar respectively. All atom simulations were carried out for 300 ns with a 2 fs timestep after minimizing and equilibrating for 20 ns. Each simulation was carried in triplicate. Trajectories were analyzed using VMD[8] and MATLAB.

*C.elegans*. Two *C.elegans* strains EAK 102 expressing htt-513(Q15) and EAK 103 expressing htt-513(Q128)[11] were obtained from Caenorhabditis Genetics Center. Worms were maintained at

20 °C in NGM plates inoculated with OP50 under standard techniques. Age synchronized worms were obtained using previously described methods[12]. EAK 102 (Q15) serves only as a control in all experiments while EAK 103 (Q128) was treated with either Ro or Benzamil.

Survival assays were performed in 96 well plates. The culture medium was prepared as described previously[12]. Final OP50 concentration in the well was set to 0.05 mg/mL. Fluorodeoxyuridine (FUdR) was used to prevent the synchronous population from reproducing[13] and the final FUdR concentration was set to 400 µM. Ro or Benzamil were added to the culture medium to obtain desired final concentration. For each well ~ 10 age synchronized L4 worms were introduced. Viability of the animals was determined every other day for up to 25 days, animals that do not move upon shaking for 30 s on a plate shaker were scored as deceased. Each condition was tested for a minimum of 50 animals per trial.

For thrashing assays, drug induced NGM plates (35 mm diameter) were prepared with a final FUdR concentration of 400 µM. For each plate ~100 age synchronized L4 worms were introduced and were maintained at 20 °C. Thrashing rate was observed on day 2 and 7 of the adulthood. Animals were picked from plates in 25 µL drops of M9 buffer and placed on a microscopic slide. They were given 30 s recovery time, and a 1 min video of their thrashing in liquid was recorded. The videos were processed and analyzed using MATLAB image processing toolbox and Image J equipped with worm tracker plugin[14,15]. A minimum of 35 animals were assessed for each condition per trial.

To measure inclusion formation via fluorescent microscopy, adult day 7 animals were obtained from NGM plates and were suspended in 15 mg/ mL 2,3-Butanedione monoxime (BDM) for 30-60 mins to induce paralysis in worms. Once paralyzed the worms were imaged in solution with either 10x magnification or 20x magnification. Inclusion counts in each image was determined by counting the fluorescent foci in each worm.

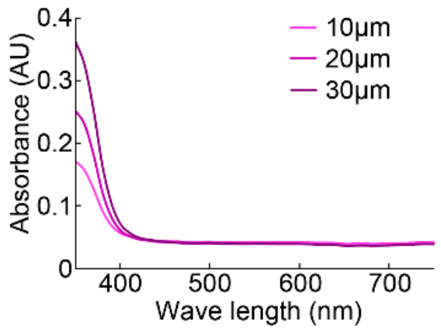


**Extended Data Figure 1**. Absorbance spectrum of Ro at various concentrations.

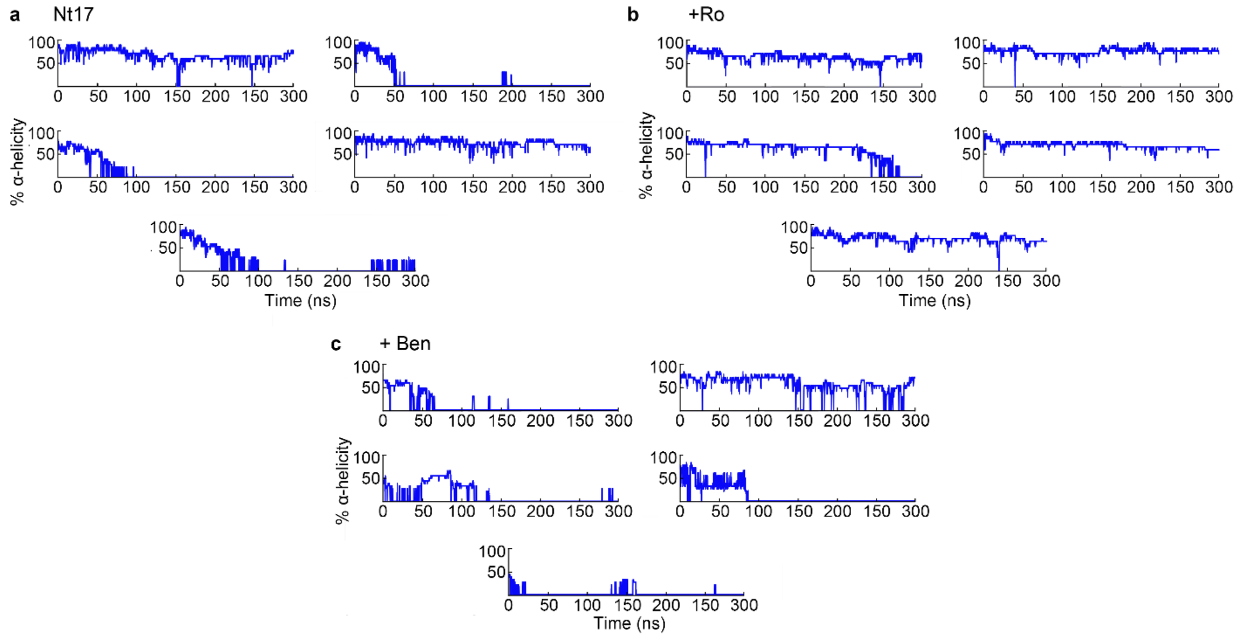


**Extended Data Figure 2**. Percentage alpha helicity of **a,** Nt17 monomer and **b,** Nt17 monomer +Ro or **c,** +Ben for five additional simulations.

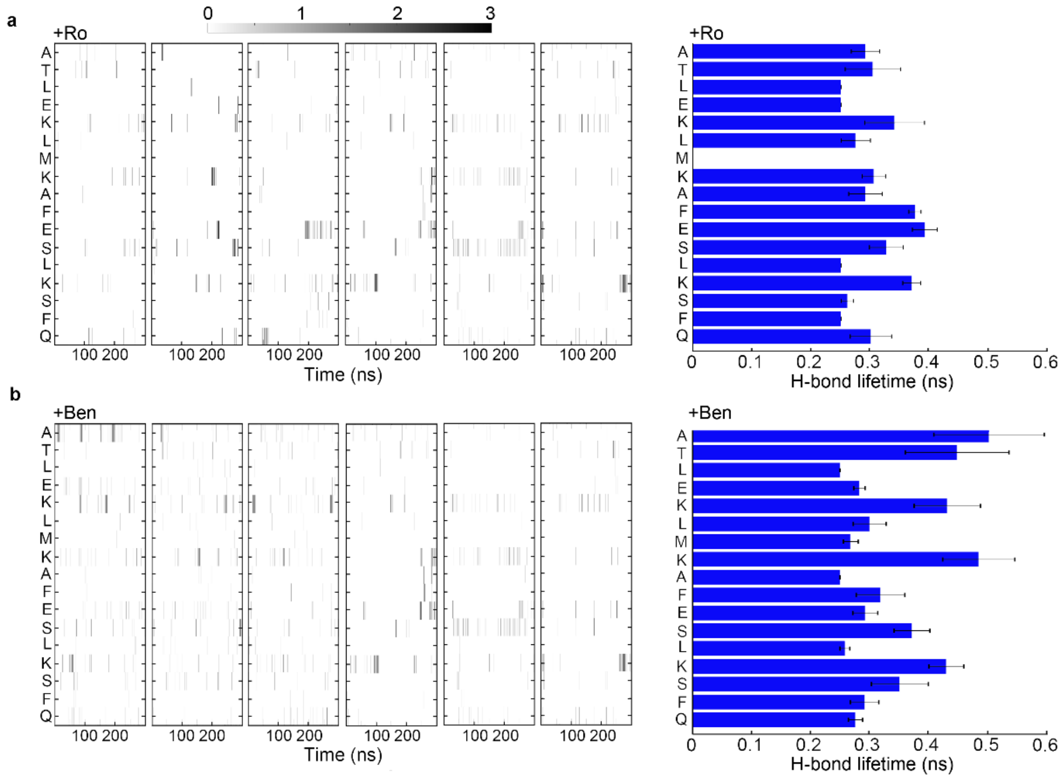


**Extended Data Figure 3.** Residue specific, time-dependent hydrogen bonding profile and average hydrogen bond lifetime of **a,** Ro and **b,** Ben with an Nt17 monomer for six independent simulations. The hydrogen bond lifetime is the mean for all occurrences across all six simulations. Error bars represent SEM.

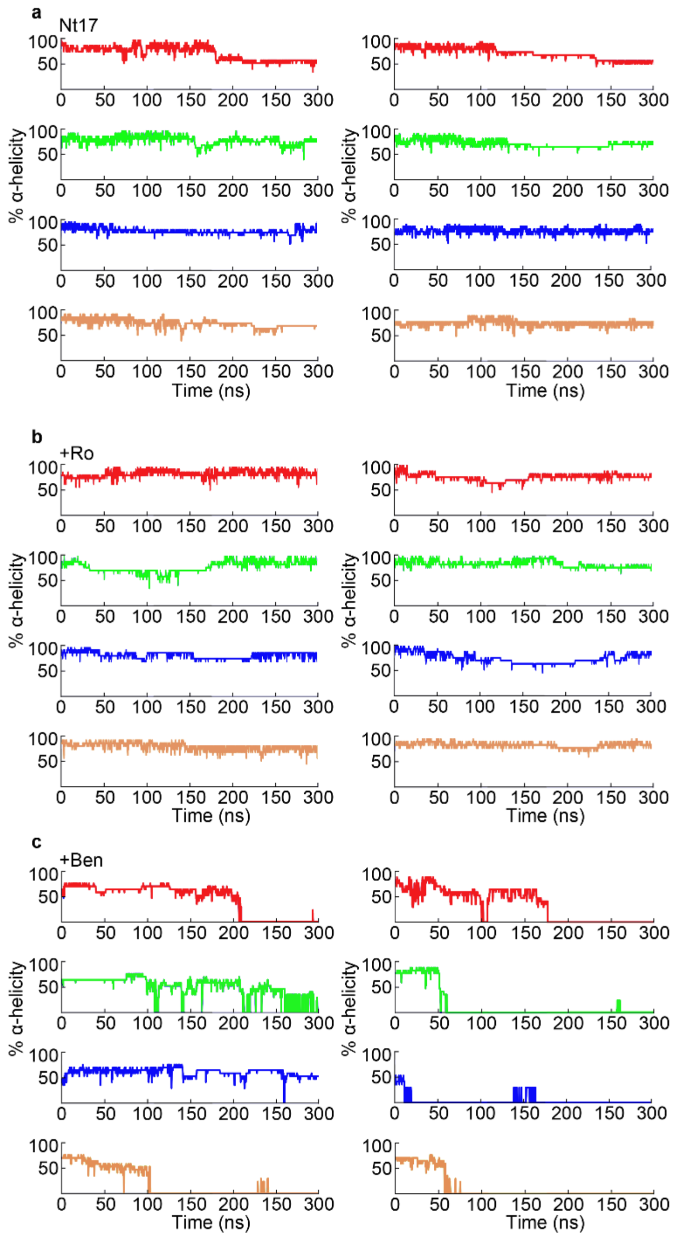


**Extended Data Figure 4.** Percent α-helicity of each peptide in the **a,** Nt17 tetramer and **b,** Nt17 tetramer +Ro or **c,** +Ben for two additional simulations.

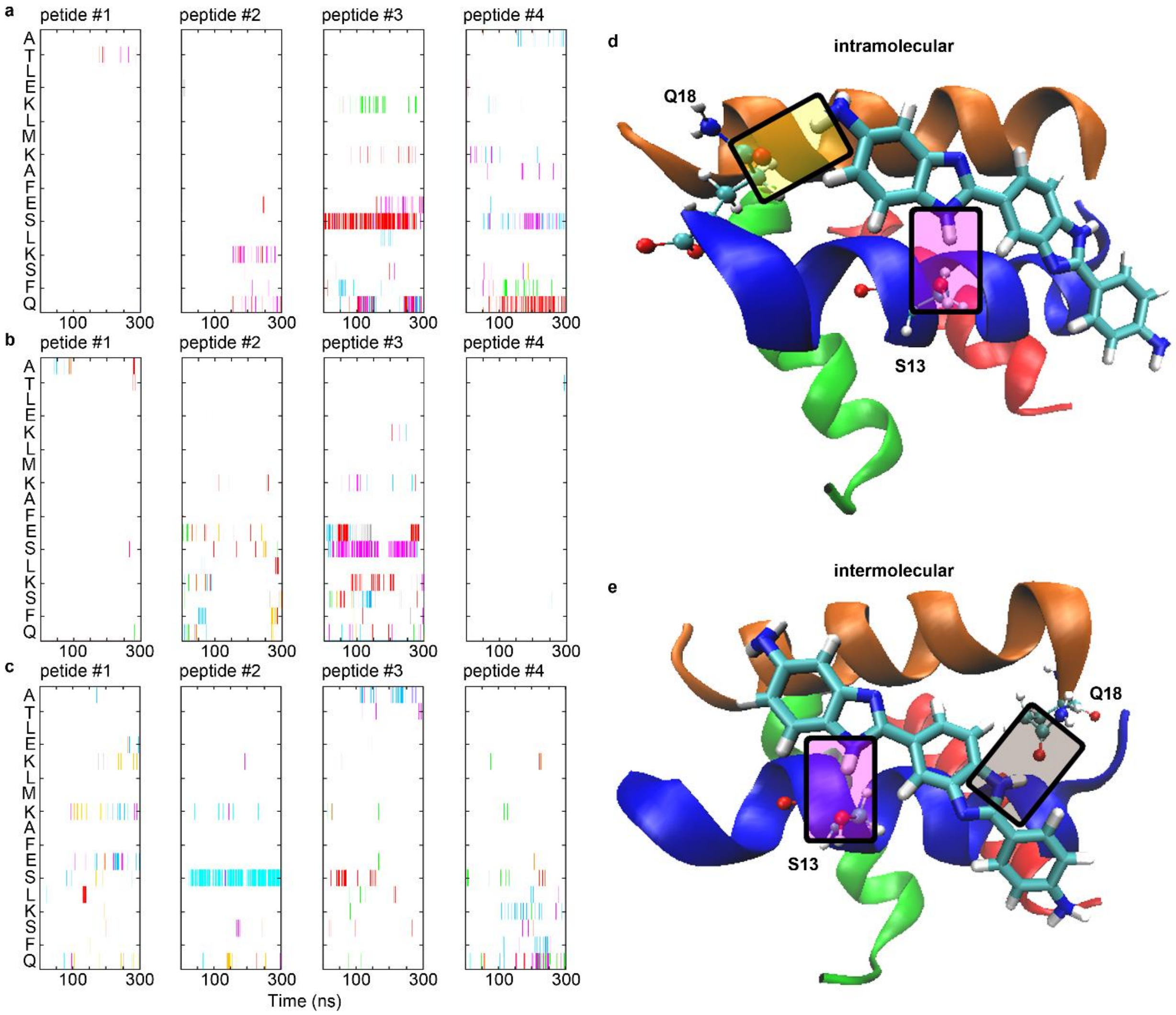


**Extended Data Figure 5**. Hydrogen bonding profile of individual peptides within the Nt17 tetramer with Ro molecules. Each color represents an individual Ro molecule. **a,b** and **c** are three independent simulations. **d**, An example of an individual Ro molecule forming hydrogen bonds with S13 and Q18 within a single peptide (intramolecular). **e**, An example of an individual Ro molecule forming hydrogen bonds with S13 and Q18 from different peptides (intermolecular). For both examples, shaded boxes indicate the hydrogen bonds.

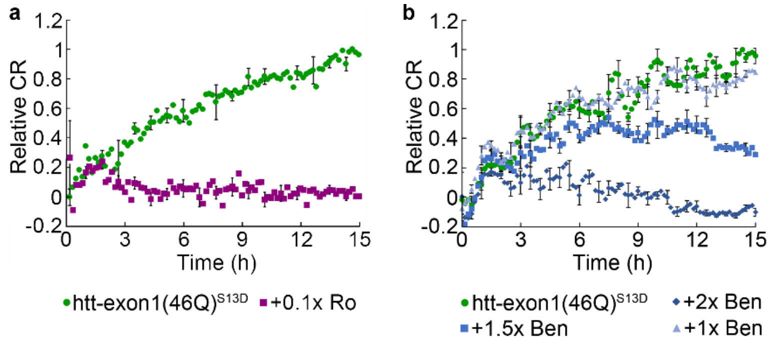


**Extended Data Figure 6**. TBLE/PDA assay of htt-exon1(46Q)$^{S13D}$. **a**, TBLE/PDA vesicles exposed to htt-exon1(46Q)$^{S13D}$ and 0.1x Ro. **b**, TBLE/PDA vesicles exposed to htt-exon1(46Q)$^{S13D}$ and various doses of Benzamil. All signals are normalized with respect to TBLE/PDA vesicle exposure to 10 µM htt-exon1(46Q)$^{S13D}$ and error bars represent SEM.
